# Partial Differential Equation (PDE) –Based Spatial Pharmacometrics in NONMEM: Method of Lines (MOL) Implementation With AI-Assisted Model Development

**DOI:** 10.64898/2026.03.01.708822

**Authors:** Yiming Cheng, Yan Li

## Abstract

Spatial heterogeneity in drug distribution, particularly within solid tumors, can compromise target engagement and drive therapeutic failure in oncology, yet it is rarely represented in population pharmacokinetic (PopPK) analyses. Standard empirical compartmental or semi-mechanistic models assume “well-stirred” tissues, and physiologically based pharmacokinetic (PBPK) models focus on organ-level distribution; neither framework directly captures intra-tumoral drug concentration gradients. Reaction–diffusion PDEs provide a mechanistic representation of penetration, spatial gradients, and washout, but routine implementation in NONMEM has been limited by operational complexity in the pharmacometrics field: native numerical templates (e.g., DOPDE/DOEXPAND-style expansions offered by NONMEM) remain cumbersome, and manual MOL coding quickly becomes labor-intensive and error-prone as geometry, initial and boundary conditions, and grid resolution change. With the emergence of advanced AI technologies, this operational barrier can be substantially reduced. This work presents a streamlined workflow for implementing spatial PDEs in NONMEM via AI tools. Using AI-assisted explicit code generation, we show how continuous spatial models can be systematically translated into coupled ordinary differential equation (ODE) systems that are directly executable in NONMEM, while keeping the resulting $DES block implementation transparent and reviewable. We illustrate the approach with one-dimensional, spherical, and two-dimensional rectangular reaction–diffusion models and provide practical guidance for iterative refinement across discretization and boundary-condition settings using appropriate prompt engineering with AI. Although AI does not reduce stiffness, remove numerical constraints, or resolve identifiability limitations, it reduces the engineering and maintenance burden of large MOL systems. When coupled with disciplined verification and appropriate scientific restraint, AI-assisted code generation can make PDE-based spatial pharmacometrics in NONMEM practical, transparent, and maintainable—supporting wider adoption of spatial modeling to interrogate target-site exposure and penetration-driven efficacy.

## Introduction

Conventional pharmacometric workflows typically infer tissue exposure from plasma pharmacokinetics (PK) using compartmental population PK (PopPK) models. ^1^ This approach is practical and widely adopted, but it implicitly treats tissues as well-stirred compartments and therefore summarizes exposure as a spatial average, effectively assuming that drug molecules crossing the vascular endothelium become instantaneously available to every cell within the tumor mass. ^1, 2^ In reality, in diffusion-limited or structurally heterogeneous tissues, most notably solid tumors, clinically meaningful concentration gradients can arise due to limited penetration, binding and consumption within tissue, and time-varying wash-in/wash-out dynamics. ^3–7^ When such gradients drive target engagement and response, spatially averaged tissue representations can obscure mechanisms that are central to efficacy and therapeutic failure. ^8^

Reaction–diffusion partial differential equations (PDEs) provide a mechanistic framework to represent penetration lag, spatial gradients, and local elimination or degradation within tissue. ^4, 7, 9, 10^ Despite their conceptual appeal, PDE-based spatial pharmacometrics has not been routinely adopted within the standard nonlinear mixed-effects ecosystem. The practical barrier is largely operational rather than theoretical: most pharmacometric platforms, including NONMEM, are optimized for ordinary differential equation (ODE) systems, whereas spatial PDEs require numerical discretization before they can be executed and maintained in production workflows. ^11^ Although PDE solution methods are mature in biomedical engineering and are often implemented in dedicated numerical environments (e.g., MATLAB), ^12^ these tools generally lack the population modeling and mixed-effects inference capabilities that are central to pharmacometrics.

In NONMEM, PDE implementation typically relies on the MOL, in which the spatial domain is discretized into layers or nodes and the PDE is converted into a coupled ODE system solvable by NONMEM’s stiff integrators (e.g., ADVAN13/LSODA). ^11^ While MOL is numerically standard, the implementation burden increases rapidly with grid size, geometry, and boundary-condition complexity: even moderate discretizations can translate into large, index-sensitive $DES blocks that are time-consuming to author, error-prone to maintain, and difficult to review. For example, manually coding N = 50 or N = 100 coupled differential equations is tedious, error-prone, and daunting. NONMEM utilities such as DOEXPAND and DOPDE can reduce some manual work; however, these workflows often rely on complex numerical “stencils” (e.g., dss044, dss020) and opaque configuration parameters (e.g., OFFSET, MVAR) that can function as a “black box” to many pharmacometricians. As a result, the capability exists, but the methodology has not been widely adopted by the broader community.

With the emergence of modern generative AI, ^13, 14^ this operational barrier can be substantially reduced. The goal of this work is to lower the barrier for PDE-based spatial pharmacometrics in NONMEM by making MOL implementations faster to generate, easier to audit, and simpler to maintain. We present an AI-assisted workflow that drafts explicit, indexed NONMEM $DES code for MOL discretizations under user-specified geometry, grid resolution, boundary conditions, and finite-difference stencil order, coupled with disciplined verification and cross-checking. We demonstrate the approach using three representative geometries commonly encountered in tissue-penetration problems: a one-dimensional slab, a spherical domain, and a two-dimensional rectangular sheet. In addition, we illustrate how grid resolution and stencil order influence MOL approximations in these settings and provide a practical prompt-design checklist to support reproducible AI-assisted generation of NONMEM-ready control streams.

Importantly, the intent of AI assistance in this context is not to change numerical stiffness, eliminate numerical constraints, or resolve fundamental identifiability limitations. Rather, AI is used as an engineering accelerator to reduce repetitive coding and formatting effort while keeping the final implementation transparent, reviewable, and aligned with established numerical methods. By focusing on explicit code generation plus verification, this workflow aims to make spatial modeling in NONMEM more accessible to pharmacometricians who wish to move beyond “average” tissue concentrations and explore penetration-driven behavior within structured tissues.

## Methods

### Software, AI Tools and Implementation

Data preparation and post-processing were performed in RStudio (R version 4.1.3, Posit Software, PBC, Boston, MA, USA). Population pharmacokinetic (PopPK) model implementation and simulation were conducted using the nonlinear mixed-effects modeling software NONMEM® (Version 7.5.1; ICON Development Solutions, North Wales, PA, USA). AI-assisted code generation was performed using Google Gemini 3.0 (Google LLC, Mountain View, CA, USA) to draft NONMEM control streams, generating explicit **$DES** blocks for method-of-lines (MOL) discretizations under user-specified geometry, grid resolution, boundary conditions, and finite-difference stencil order. Prompts were designed to request (i) syntactically valid/correct NONMEM code and (ii) transparent, explicitly indexed differential equations suitable for direct execution in ADVAN13. All AI-generated control streams were reviewed for syntactic validity and subjected to manual spot checks of indexing, stencil coefficients, and boundary-condition logic. In addition, code generation outputs were independently cross-validated using ChatGPT-5.2® (OpenAI, San Francisco, CA, USA) to reduce automation bias and to triangulate consistency of the discretization and implementation.

### MOL Implementation

The MOL was used to implement reaction–diffusion PDE models within NONMEM by converting the continuous spatial problem into a finite-dimensional system of ordinary differential equations (ODEs). Briefly, the spatial domain was discretized into a fixed grid of N layers (1D slab or spherical radius) or N_x_ * N_y_ nodes (2D rectangular sheet) with spacing (Δx and Δy for 2D). Spatial derivatives were approximated using symmetric finite-difference stencils, in which the spatial second derivative at a given grid point is expressed as a weighted combination of concentrations at neighboring points. Specifically, we used either a three-point stencil (center node plus its immediate neighbors; second-order accurate) or a five-point stencil (center node plus two neighbors on each side; fourth-order accurate), depending on the scenario and sensitivity analyses. This discretization converts the PDE operator into algebraic couplings between neighboring grid nodes, which were implemented explicitly in the NONMEM **$DES** block. Boundary conditions were encoded directly in $DES by modifying the update equations at boundary nodes, using flux-matched conditions at tissue interfaces and no-flux (Neumann) or symmetry conditions at distal boundaries (e.g., ∂𝐶/ ∂𝑥 ∣_𝑥=𝐿_= 0 in slab geometry and the appropriate limiting form at r = 0 in spherical geometry). For the 2D sheet, a row-major mapping was used to translate (x, y) grid coordinates into the 1D state vector required by NONMEM, and neighbor connectivity in the x and y directions was implemented using index offsets of ± 1 and ± N_x_, respectively. Grid resolution (N or N_x_, N_y_) and stencil order were varied in dedicated sensitivity analyses to assess discretization dependence, with higher-resolution settings used as numerical reference solutions for comparison.

### The One-Dimensional Spatiotemporal Drug Distribution Model

The one-dimensional spatiotemporal drug distribution model comprises a central plasma compartment (A1) with one-compartment IV disposition, coupled to a spatially discretized tissue slab representing drug penetration into tissue. The plasma compartment serves as the driving reservoir: drug is eliminated via systemic clearance and exchanges with the tissue surface through a discretization-derived diffusive coupling. For interpretability, the plasma–surface coupling can be written in concentration form as:

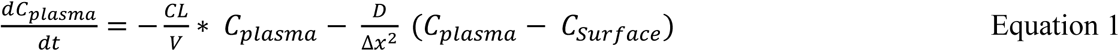

Within the tissue slab, concentrations evolve according to the reaction–diffusion PDE:

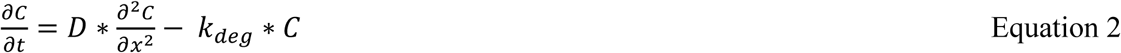

where CL is plasma clearance, V is the central volume of distribution, C_plasma_ is the plasma drug concentration, C_surface_ is the tissue-surface concentration at the plasma–tissue interface, *D* is the diffusion coefficient, Δx is the spatial grid spacing, k_deg_ is the first-order degradation rate within tissue, Δx is L/(N-1), where L is slab thickness and N is the number of layers. Note that for this illustrative model, the plasma-tissue coupling term in Equation 1 is presented in a simplified concentration-driven form; explicit geometric scaling factors (e.g., interfacial surface area relative to plasma volume) are treated as effectively absorbed into the transport term or normalized to unity to maintain focus on the numerical discretization workflow.

### Spherical Reaction–Diffusion Spatiotemporal Drug Distribution Model

The spherical reaction–diffusion model extends the one-dimensional slab framework by replacing the linear tissue domain with a spatially discretized spherical tumor (0 < r <R). The central plasma compartment (A1) and its dynamics are identical to those in the 1D slab model (Eq. 1), and the plasma compartment is coupled to the tumor surface through a discretization-derived diffusive coupling at r = R.

Within the spherical tissue, drug concentrations evolve according to the reaction–diffusion PDE in radial coordinates:

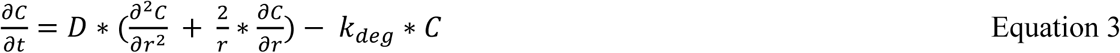

where D is the diffusion coefficient, r is the radial distance from the tumor center, and k_deg_ is the first-order degradation rate within tissue.

### 2D Rectangular Reaction–Diffusion Spatiotemporal Drug Distribution Model

The 2D rectangular reaction–diffusion toy example extends the 1D slab and spherical frameworks to a two-dimensional tissue sheet discretized on an N_x_ * N_y_ grid (N_x_ = N_y_ = 10). The systemic plasma compartment (A1) followed one-compartment IV disposition as in Eq. 1. Tissue concentrations evolve according to the 2D reaction–diffusion PDE:

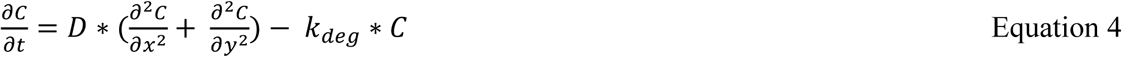

where D is the diffusion coefficient and 𝑘_deg_(𝑥, 𝑦)is a first-order degradation rate. For implementation in NONMEM, a row-major mapping was used to translate (x, y) grid coordinates into the 1D state vector required by NONMEM, with neighbor connectivity implemented through index offsets of ±1(in the 𝑥-direction) and ±𝑁_𝑥_(in the 𝑦-direction).

## Results

### Characterization of One-Dimensional Spatiotemporal Drug Distribution using the MOL With AI-Assisted Model Development

In the first toy example, we generated a ready-to-run NONMEM control stream that implements a one-dimensional reaction–diffusion PDE using the numerical MOL (Supplementary NONMEM code #1). The model describes drug transport within a linear tissue slab (0 < x < L) as defined in Equations 1 and 2. The primary purpose of this example was to evaluate whether an AI-assisted workflow can reliably produce the large, index-sensitive NONMEM $DES code required for an explicit MOL implementation, and to document the verification steps used to confirm numerical correctness.

Using Gemini 3 as a code-generation assistant, we generated a syntactically valid NONMEM $DES block containing explicit differential equations for a 50-layer spatial discretization (N = 50). The prompt specified a one-dimensional linear (slab) geometry, a flux-matched condition at the left boundary and a no-flux (Neumann) condition at the right boundary, a fourth-order five-point symmetric finite-difference stencil for the spatial second derivative, and an N = 50 grid specification; we subsequently verified that the generated equations implemented the intended stencil across interior nodes. This automated generation replaced manual hand-coding of the coupled ODE system, which is typically error-prone due to repetitive indexing and boundary-transition logic.

To move beyond syntax validation and ensure that the AI-generated code correctly instantiated the intended MOL discretization, we performed a structured verification consisting of three checks. First, we verified that the interior stencil matched the theoretical five-point formulation for the Laplacian, including coefficient symmetry and index placement (e.g., −𝐴_𝑖−2_ + 16𝐴_𝑖−1_ − 30𝐴_𝑖_ + 16𝐴_𝑖+1_ − 𝐴_𝑖+2_). Second, we confirmed that the interior stencil was applied consistently across the full interior region (Compartments 4–49), and that the transition logic near the boundaries did not introduce off-by-one indexing or missing neighbor terms. Third, we reviewed the boundary conditions implemented in the **$DES** block to confirm that they matched the intended physical assumptions: (i) a flux-matched condition at the plasma–tissue interface (x = 0) and (ii) a Neumann no-flux condition at the distal boundary 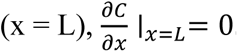. No indexing or boundary-transition errors were identified during these checks.

Figure 1 presents the simulated spatiotemporal drug distribution, and Table 1 summarizes the simulation parameters. The results show a steep early-time spatial gradient, indicating a diffusion-driven lag in penetration toward deeper layers. Over time, the gradient relaxes as the drug distributes across the slab. As plasma concentrations decline, net flux can reverse (tissue-to-plasma back-diffusion), producing a washout phase governed by the combined effects of back-transport and intratissue degradation (k_deg_).

**Figure 1.**
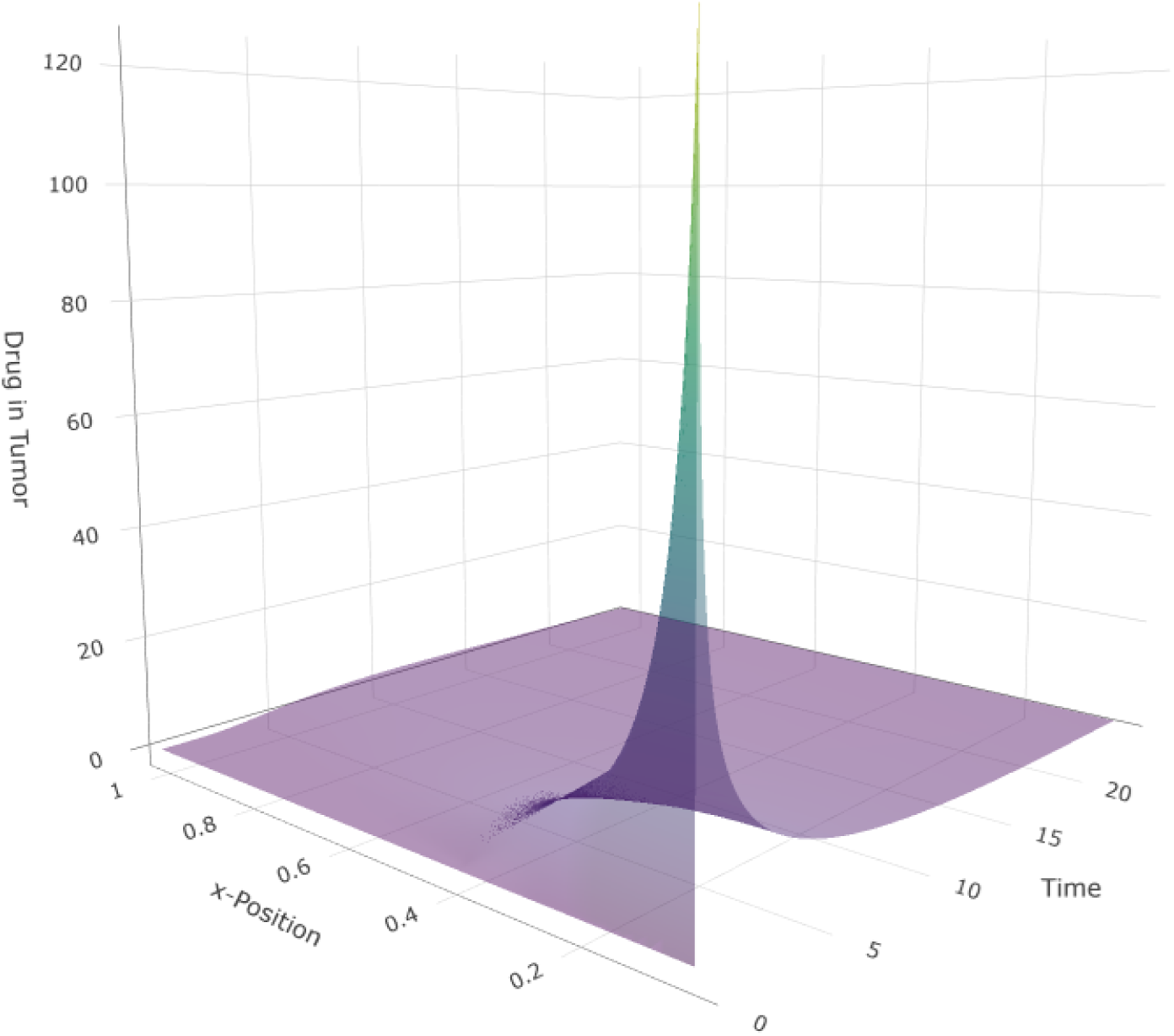
Simulated spatiotemporal drug distribution from the one-dimensional reaction–diffusion model.

**Table 1.** Parameters used in the one-dimensional, spherical, and two-dimensional rectangular reaction–diffusion models.

| **PK parameter** | **Values** |
| --- | --- |
| **CL (L/hr)** | **5.0** |
| **V (L)** | **36.0** |
| **Diff (cm<sup>2</sup>/hr)** | **0.02** |
| **Kdeg (1/hr)** | **0.01** |

Abbreviations: CL, plasma clearance; Diff, diffusion coefficient; Kdeg, first-order degradation rate within tissue; V, central compartment volume.

### Discretization Sensitivity and Rapid Code Regeneration Using AI-Assisted MOL Implementations

Given that both the finite-difference stencil (three-point vs. five-point symmetric schemes) and the spatial grid resolution (number of layers, N) directly govern the numerical accuracy of a method-of-lines (MOL) approximation, we evaluated the sensitivity of simulated tissue exposure to these two discretization choices. This analysis also served as a practical test of how flexibly the AI-assisted workflow could regenerate NONMEM-ready code under alternative discretization specifications.

Specifically, we prompted Gemini 3 to generate updated NONMEM control streams for a factorial set of scenarios combining (i) grid sizes N = 10, 25, 50 and 100 layers and (ii) either a three-point or five-point symmetric finite-difference stencil for the spatial second derivative. As a numerical reference, we used the solution obtained with N = 100 and the five-point stencil as the “high-resolution” benchmark, and quantified deviations in predicted tissue (tumor) concentrations for each alternative discretization.

For each scenario, Gemini 3 produced the corresponding revised **$DES** code rapidly and without syntactic issues, eliminating substantial manual re-coding effort associated with re-indexing and boundary-condition bookkeeping when N and stencil order change.

Figure 2A compares results for N = 50 under the three-point versus five-point symmetric stencil. Under the current parameterization and time horizon, the two stencils yielded nearly indistinguishable spatiotemporal tissue profiles, indicating that the lower-order three-point stencil provided adequate accuracy for subsequent analyses. Accordingly, all remaining simulations in this work used the three-point symmetric stencil for computational simplicity.

**Figure 2.**
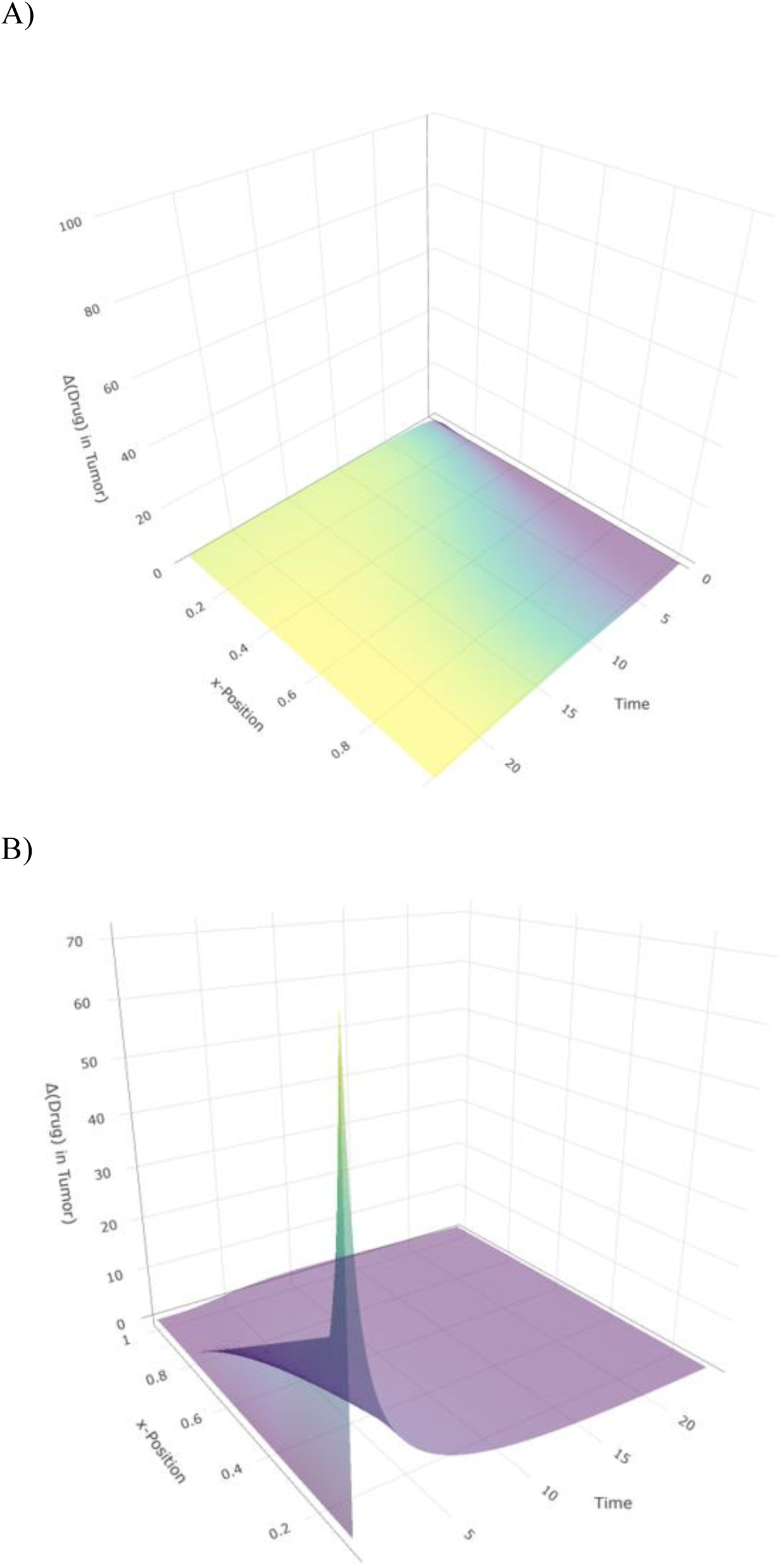
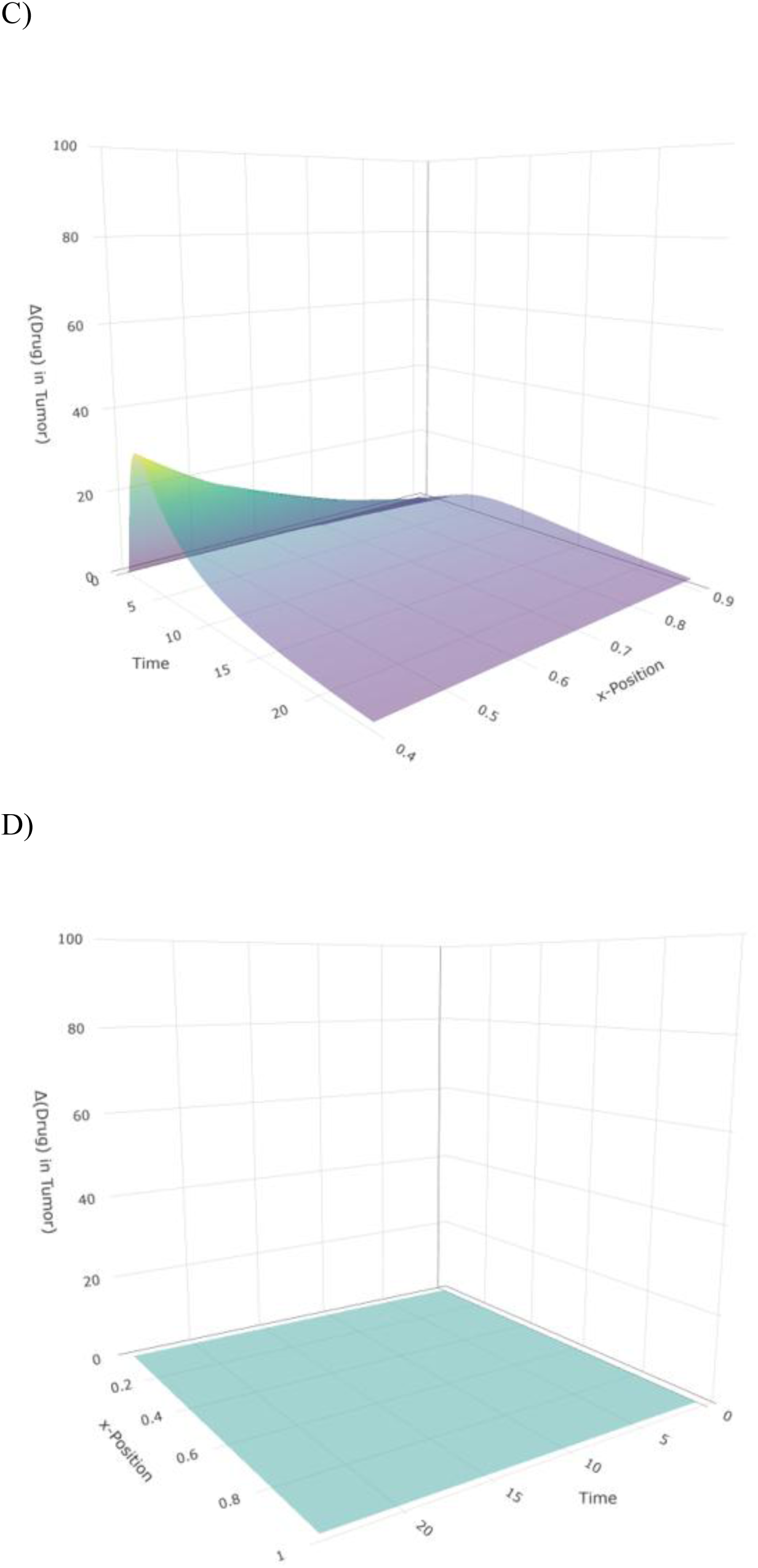
Sensitivity of the method-of-lines (MOL) approximation to grid resolution and finite-difference stencil order. (A) Comparison at N = 50 using three-point versus five-point symmetric stencils. (B–D) Comparison of grid resolutions N = 10, N = 25, and N = 50, respectively, against the N = 100 reference solution.

Figures 2B–2D compare grid resolutions (N = 10, 25, 50) against the N = 100 benchmark. The N = 50 solution closely matched the N = 100, whereas coarser grids (N = 10 and N = 25) showed clear deviations in predicted tissue concentrations, consistent with discretization error from insufficient spatial resolution. These results highlight the importance of routinely assessing grid dependence in MOL-based spatial models; overly coarse discretizations can materially bias predicted target-site exposure even when the underlying PDE specification is unchanged.

### AI-Assisted MOL Implementation of Spherical Reaction–Diffusion Spatial Models in NONMEM

In the second toy example, we extended the MOL implementation from a linear slab to spherical tumor geometry (0 < r <R), where the governing operator includes the spherical Laplacian and introduces layer-specific geometric terms, as defined in Equations 1 and 3 (Supplementary NONMEM code #2).

Using Gemini 3 as a code-generation assistant, we generated a syntactically valid NONMEM $DES block for a 50-layer radial discretization (N = 50). The prompt specified spherical geometry, a flux-matched boundary at the tumor surface (r = R), a symmetry condition at the tumor core (r = 0), and a three-point symmetric finite-difference discretization adapted to the spherical operator. Compared with the slab case, spherical MOL implementations are substantially more error-prone to code manually because the geometric coefficient multiplying the radial-gradient term must be computed and applied uniquely for each layer (i.e., the coefficient changes with r at every grid point).

To confirm correctness beyond syntax, we performed three targeted verification checks tailored to spherical coordinates. First, we confirmed that the generated equations implemented the expected spherical operator structure, including both the second-derivative term 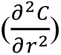 and the geometry-dependent gradient term 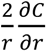, with appropriate layer-wise coefficient application. Second, we spot-checked selected layer indices to verify that the precomputed geometric coefficients corresponded to the physical radii of those grid points (i.e., the coefficient magnitude scaled appropriately with depth). Third, we verified the r = 0 boundary handling, confirming that the core equation used the appropriate limiting form for the spherical Laplacian at the origin (via the standard L’Hôpital-type treatment), including the required multiplicative factor in the center-node update. No indexing, boundary, or geometric-coefficient inconsistencies were identified in these checks.

Figure 3 visualizes the resulting radial concentration gradients over time. Early after dosing, concentrations are highest near the tumor surface, reflecting the boundary influx and limited early-time penetration. By intermediate times (e.g., t = 6 to t = 12), the profiles show progressive inward diffusion, with the concentration gradient flattening as drug distributes toward the core. At later times (e.g., t = 18 to t = 24), overall concentrations decline across all radii, consistent with systemic decline and intratumoral loss processes, while the spatial gradient continues to diminish. Collectively, these results illustrate the characteristic penetration lag and subsequent washout behavior that cannot be represented by well-stirred tumor compartments, and they demonstrate that the AI-assisted MOL workflow extends cleanly to spherical PDE implementations in NONMEM.

**Figure 3.**
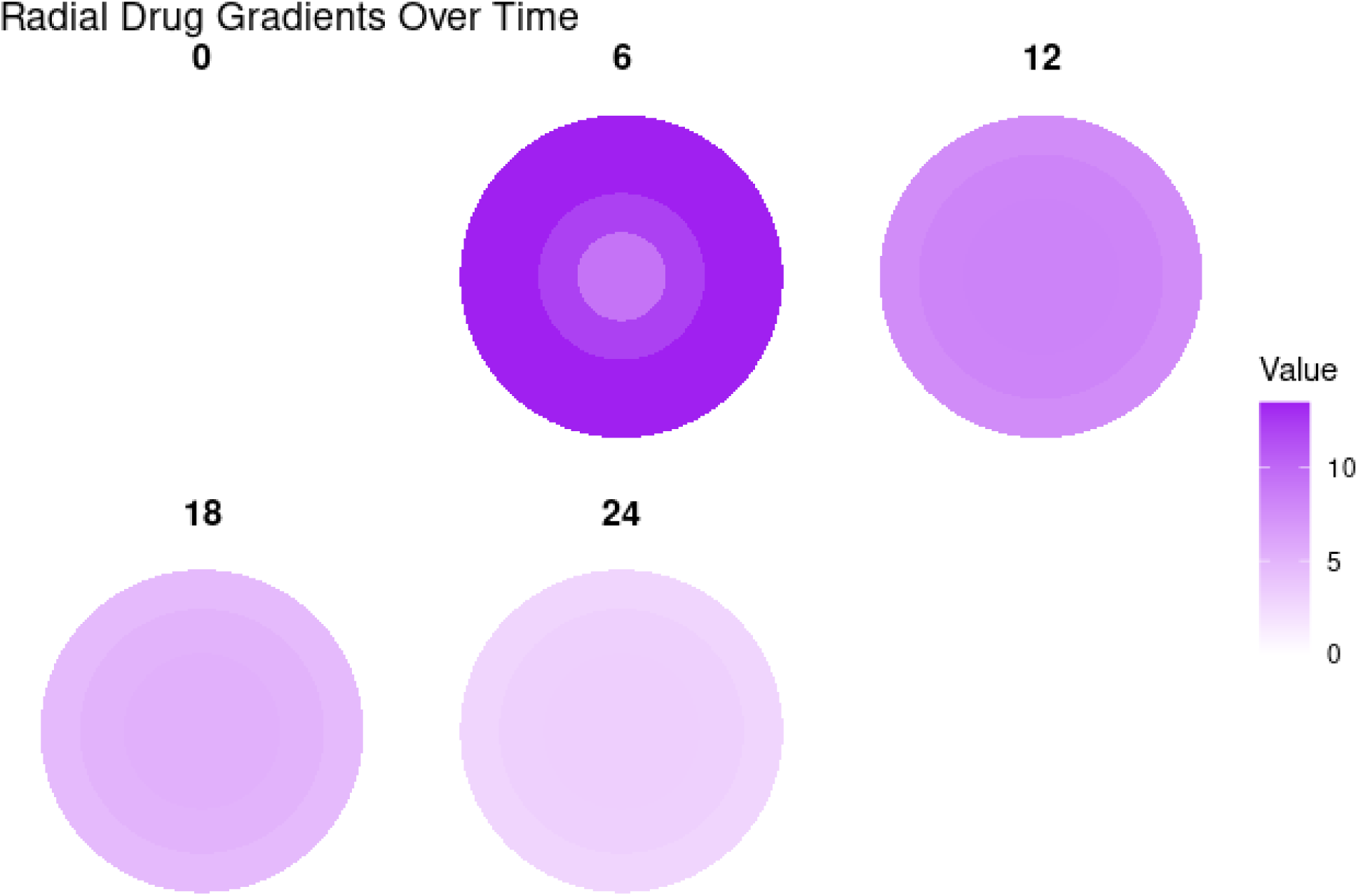
Simulated spatiotemporal drug distribution from the spherical reaction–diffusion model. Panels correspond to different time points.

### AI-Assisted 2D Rectangular-Grid Method-of-Lines Implementation in NONMEM

In the third toy example, we implemented a two-dimensional reaction–diffusion PDE in NONMEM using the MOL on a rectangular grid. The model represents drug transport within a 10-by-10 tissue sheet, where diffusion occurs in both the x- and y-directions using a single diffusion coefficient, as defined in Equations 1 and 4 (Supplementary NONMEM code #3). Because the grid spacing was identical in both directions (Δx = Δy), and the same diffusion coefficient D is used in both directions, the discretization is isotropic.

Using Gemini 3 as a code-generation assistant, we generated a syntactically valid NONMEM **$DES** block containing explicit differential equations for a 100-compartment spatial discretization (N = 100), plus a central plasma compartment. The prompt defined an intentionally heterogeneous forcing structure: plasma exchange was restricted to the left boundary (x = 0) of the central source rows, while accelerated loss was imposed at the periphery by increasing the degradation rate in the top and bottom rows. The plasma reservoir was transient (IV bolus–like), with systemic elimination from the central compartment, such that the source strength decayed over time.

We verified the generated code with targeted checks focused on common failure modes in 2D MOL implementations. First, we confirmed that interior nodes implemented the expected five-point 2D stencil (center plus four neighbors), with correct horizontal coupling via indices i ± 1 and vertical coupling via indices i ± N_x_, and without erroneous wrap-around across row boundaries. Second, we verified that the plasma-coupling (“source”) terms were confined to the specified central rows. Third, we confirmed that the high-degradation (“sink”) specification was applied only to the intended boundary rows and the adjacent row, with all remaining interior rows retaining the baseline degradation rate. No indexing, connectivity, or parameter-assignment inconsistencies were identified in these checks.

Figure 4 visualizes the resulting spatiotemporal concentration field. At early time points, drug accumulates preferentially within the central source band, producing a localized peak centered in the mid-𝑦 region. As time progresses, diffusion spreads drug laterally and vertically, broadening the distribution while the peak magnitude diminishes due to the declining plasma reservoir and ongoing degradation. At intermediate times (e.g., t = 12 and t = 18), the field becomes progressively flatter as the source weakens, with more rapid attenuation near the sink rows. By t = 24, the concentration gradient has largely dissipated, consistent with a transient source coupled with sustained peripheral sinks and bulk degradation.

**Figure 4.**
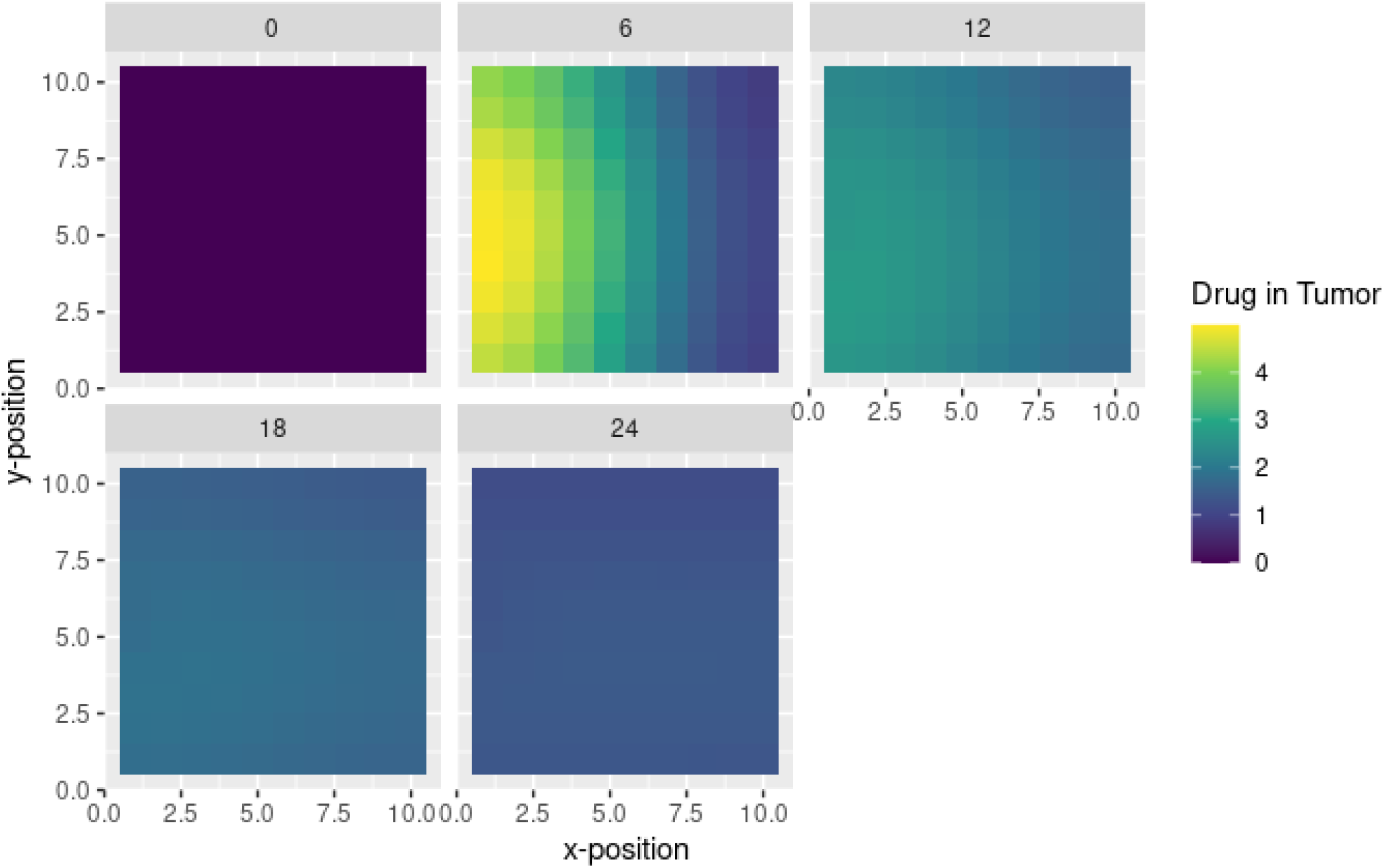
Simulated spatiotemporal drug distribution from the 2D rectangular reaction–diffusion model. Panels correspond to different time points.

## Discussion

A limitation of conventional pharmacometric workflows is that plasma PK is often an imperfect surrogate for target-site exposure, particularly in diffusion-limited or heterogeneous tissues such as solid tumors. ^3, 15–17^ Standard compartmental PopPK models treat tissues as well-stirred and therefore predict an average concentration—whether in empirical multi-compartment models or in many semi-mechanistic structures—which can mask clinically meaningful spatial gradients. ^1, 2^ In contrast, reaction–diffusion PDEs can represent penetration lag, spatial gradients, and washout behavior that are central to understanding target engagement and therapeutic failure in solid tumors and other structured tissues. ^5, 18–22^ The barrier to broader adoption has not been conceptual, but operational: spatial PDEs have been difficult to implement within the population modeling ecosystem routinely used by pharmacometricians.

NONMEM is fundamentally an ODE-based platform and is not designed to directly solve PDEs. ^11^ Implementing PDEs in NONMEM therefore typically relies on the MOL, which discretizes space into multiple layers and approximates the PDE as a coupled system of ODEs solvable by NONMEM’s stiff integrators (e.g., ADVAN13/LSODA). ^11^ Although MOL is numerically standard, the implementation burden increases rapidly: even moderate grids translate into hundreds of ODE statements, together with extensive indexing initial- and boundary-condition logic. This creates a high risk of implementation errors (e.g., off-by-one indexing, incorrect neighbor coupling, wrap-around at boundaries, or incorrect coefficients). Such errors can still yield numerically stable runs and plausible-looking profiles, while materially violating the intended physics. Since NONMEM 7.4, utility programs such as DOEXPAND and DOPDE have been available to reduce the amount of manual effort required. ^11^ In practice, adoption has remained limited because these workflows can require complex configuration and can be difficult to debug when large systems are generated. Consistent with this, PDE/MOL implementations in NONMEM remain relatively uncommon in the pharmacometrics literature.

With the emergence of modern generative AI, this operational barrier can be substantially reduced. ^13, 14^ The key value of AI in this context is not that it changes the theory—MOL and finite-difference PDE discretization are mature—but that it removes a practical bottleneck: the time-consuming, error-prone task of authoring and refactoring large, index-sensitive **$DES** blocks. This paper provides a practical path toward routine PDE-based spatial modeling in NONMEM by combining explicit MOL implementations with AI-assisted code generation and a verification mindset that makes large ODE systems feasible to author, review, and iterate. The primary contribution is therefore not a new numerical method, but a reproducible workflow that makes spatial PDE modeling accessible without requiring deep familiarity with pattern-based PDE machinery.

In the first toy example, AI-assisted code generation produced a NONMEM-ready MOL implementation of a one-dimensional reaction–diffusion model. The simulated profiles reproduced the expected qualitative behavior: an early steep spatial gradient reflecting diffusion-limited penetration (penetration lag), followed by progressive gradient relaxation, and eventual washout as the plasma reservoir declined and intratissue degradation/back-diffusion dominated. Because AI can rapidly regenerate code under alternative discretization settings, we also evaluated discretization sensitivity across grid resolution (N = 10, 25, 50 and 100) and symmetric stencil order (three-point vs five-point). These experiments showed that grid resolution was the dominant driver of differences: N = 50 closely matched the high-resolution benchmark, whereas coarser grids deviated. At N = 50, three- and five-point stencils produced similar profiles under the tested conditions, supporting use of the simpler stencil in subsequent analyses and reinforcing the importance of routine grid-dependence checks.

We then extended the workflow to more complex geometries, where manual coding is typically daunting and more error-prone. In the spherical tumor example, correct implementation requires layer-dependent geometric coefficients and careful handling of the r = 0 singularity. The AI-assisted workflow successfully generated the required equations and enabled explicit verification of the core symmetry/limiting form. In the two-dimensional rectangular grid example, the major implementation risk is correct 2D-to-1D indexing and neighbor connectivity in both directions without wrap-around errors at row boundaries. The generated code correctly instantiated the 2D stencil and supported spatially heterogeneous source–sink structures, producing the expected spreading-and-decay concentration patterns over time.

Collectively, these examples illustrate that AI does not reduce stiffness, eliminate numerical constraints, or resolve identifiability limitations. What it changes is the engineering and maintenance burden of large MOL systems. In this work, AI-assisted generation enabled rapid creation of NONMEM-ready **$DES** code across multiple geometries and discretization settings, substantially lowering the time and error risk traditionally associated with MOL implementations. This capability is particularly valuable because spatial modeling should be iterative by design: discretization, boundary assumptions, and geometry are often refined as the scientific question and available data evolve. To facilitate adoption, we summarize a prompt-design checklist for AI-assisted generation of MOL ODEs and NONMEM control streams in Table 2.

**Table 2.** Prompt design checklist for AI-assisted generation of method-of-lines ODEs and NONMEM code.

| Checklist item | Options (examples) |
| --- | --- |
| Geometry | Linear/slab (e.g., skin patch);<br>Spherical (e.g., solid tumor);<br>Cylindrical (e.g., blood vessel). |
| Boundary conditions | For slab: $x = 0$ (inner) and $x = L$ (outer).<br>For spherical/cylindrical: $r = 0$ (inner) and $r = R$ (outer).<br>Options:<br>• Flux input (e.g., from plasma compartment);<br>• Fixed concentration (Dirichlet, e.g., $C = C_{\text{plasma}}$ or $C = 0$);<br>• No-flux/reflecting (Neumann, e.g., $\partial C / \partial x = 0$ or $\partial C / \partial r = 0$);<br>• Sink/transfer (mass-transfer/Robin, e.g., $\text{flux} = k \cdot C$ at the boundary) |
| Initial condition | Tissue concentration at $t = 0$ (e.g., all layers start at 0, or baseline steady state if applicable) |
| Grid specification | Number of layers $N$ (e.g., 20 or 50);<br>Physical length $L$ (slab) or radius $R$ (spherical/cylindrical) (e.g., 1 cm) |
| Stencil preference | 3-point (standard): simpler, robust, fewer ODE statements;<br>5-point (higher accuracy): more complex, more ODE statements |

At the same time, AI introduces a well-recognized risk: automation bias. Syntactically correct code may still encode incorrect physics. For this reason, AI-assisted PDE workflows should be paired with routine, audit-friendly verification. Based on common failure modes in MOL implementations, we recommend a minimal QC checklist whenever AI is used to generate PDE/MOL code: (1) stencil checks (spot-check multiple interior nodes for correct coefficients and neighbor indices); (2) boundary-condition checks (verify left/right or core/surface equations match intended physics such as flux, no-flux, or symmetry conditions); (3) geometry checks when applicable (validate layer-dependent coefficients and correct limiting treatment at singular points; (4) grid dependence checks (confirm stability of key conclusions as N increases); and (5) cross-validation where feasible (compare against an independent implementation in R/Python/MATLAB or triangulate with a second AI-generated discretization from the same specification).

Several limitations warrant emphasis. First, spatial parameters—especially diffusion coefficients—are often weakly identifiable from plasma data alone. Robust inference may require tissue observations, complementary biomarkers, or external information (e.g., in vitro/ex vivo experiments) that can be incorporated as informative priors or constraints during estimation. Second, computational scaling is nontrivial as grid size and dimensionality increase, which can impose memory or runtime constraints. Consistent with this, when we increased the 2D grid resolution (e.g., N_x_ = N_y_ = 20), NONMEM produced a resource-related error (“SIZE OF NMPRD4 EXCEEDED; LNP4 IS TOO SMALL, INCREASE USING $SIZES”), highlighting practical limits that must be managed via solver settings, memory sizing, or model reduction strategies. Third, the toy examples presented here are intended to establish feasibility and verification practices; application to real datasets will require careful alignment of geometry, boundary assumptions, and observation/measurement models, as well as explicit consideration of identifiability and experimental design.

Overall, this work demonstrates that spatial PDE-based modeling in NONMEM—long considered possible but operationally difficult—can become practical with AI-assisted code generation, provided that the workflow is coupled with systematic verification and appropriate scientific restraint regarding identifiability and computational limits.

## Conclusion

Spatial PDE models provide a mechanistic bridge between plasma PK and target-site exposure by representing diffusion-limited gradients that well-stirred compartments cannot capture. Although NONMEM can solve MOL-derived ODE systems, adoption has been limited by the implementation burden and error risk of large, index-sensitive equation blocks. The results in this work demonstrate that AI-assisted code generation—combined with disciplined verification—can make PDE-based spatial pharmacometrics in NONMEM practical, transparent, and maintainable, enabling wider use of spatial modeling to interrogate target-site exposure and penetration-driven efficacy.

## Consent for publication

All the authors have reviewed and concurred with the manuscript.

## Funding

This work was sponsored and funded by Bristol Myers Squibb.

## Authors’ contributions

Y.C. and Y.L. made substantial contributions to conception and design, acquisition of data, or analysis and interpretation of data; took part in drafting the article or revising it critically for important intellectual content; agreed to submit to the current journal; gave final approval of the version to be published; and agreed to be accountable for all aspects of the work.

## Notes

### Competing Interest Statement

The authors have declared no competing interest.

